# Cell-extrinsic effects of *Irf2* on cDC2 development

**DOI:** 10.1101/2023.08.24.554658

**Authors:** Kevin W. O’Connor, Tiantian Liu, Sunkyung Kim, Theresa L. Murphy, Kenneth M. Murphy

## Abstract

Characterization of the functional effects of cDC2s in vivo requires model systems in which cDC2s are depleted. Previous literature has reported a loss of cDC2s in mice lacking the transcription factor IRF2^1,2^. We sought to further characterize the cDC2 defect in these animals. Here, we find that the requirement for IRF2 in cDC2 development and survival is cell-extrinsic and correlated to the development of dermatitis in the *Irf2*^*-/-*^ model system. We also find that Flt3L-mediated in vitro development of cDC1s and cDC2s, but not pDCs, is abrogated in *Irf2*^*-/-*^ bone marrow, as well as in wild-type bone marrow cultured with IFNα. Loss of interferon α (IFNα) signaling in *Irf2*^*-/-*^ mice restored cDC2 development in vivo and cDC1 and cDC2 development in vitro. We therefore conclude that IRF2 is required for cDC2 development in a cell-extrinsic manner dependent on IFNα signaling.

## Introduction

The transcriptional basis for cDC2 development is poorly understood. In mice, cDC1 development and pDC development can both be selectively abrogated by genetic deletion of specific transcription factors or enhancers^3^. In contrast, no known transcription factor or enhancer knockout mutant can specifically and completely prevent cDC2 development. Our lab has recently developed a mouse model lacking cDC2 development owing to triple mutations in the *Zeb2* enhancer, but this mouse also lacks monocyte development, complicating analysis of cDC2 function^4^. Additionally, there are currently no genetic models that enable specific targeting of cDC2s via either Cre recombinase expression or DTR-mediated ablation^3^. Lack of a model in which cDC2s are specifically depleted has significantly hindered ability to answer key questions related to cDC2 function, such as whether cDC2s are required for rejection of tumors or formation of antiviral responses.

Several transcription factors have been identified as being important for development of specific DC2 populations or for execution of specific DC2 functions. Notch2 is required for the development of the ESAM^+^ subset of splenic Sirpα^+^ DC2s and for the development of small intestine lamina propria CD103^+^ CD11b^+^ DC2s^5^. These populations both also require signaling through the lymphotoxin β receptor (LTβR)^6,7^, and in the case of splenic ESAM^+^ DC2s, it was specifically shown that RelB is a required effector of the LTβR signaling pathway^8^. The transcription factor Klf4 is required for development of peripheral lymph node CD103^-^ CD11b^-^ Sirpα^+^ DC2s and lung Mgl2^+^ Sirpα^+^ DC2s^9^. Additionally, CD11c-cre mediated deletion of Klf4 in dendritic cells resulted in a 50% reduction in splenic DC2 numbers and a failure to induce an appropriate Th2 response to *S. mansoni*^*9*^. Lastly, IRF4 is required for DC2 migration and *IRF4*^*-/-*^ mice lack migratory DC2 populations in many lymph nodes, but DC2s are still present in non-lymphoid tissues of these mice^10^.

The transcription factor IRF2 has been reported to be important for DC2 development in the spleen^1,2^. IRF2 is a member of the Interferon Regulatory Factor family of transcription factors, of which two other members, IRF4 and IRF8, already have established roles in DC development^11^. IRF family members are characterized by the presence of a highly conserved DNA-binding domain and a more variable IRF-associating domain (IAD)^12^. The IAD region is required for IRF family transcriptional activity. Interactions between IRF8 and PU.1 drive transcriptional activity from E protein-IRF combinatorial elements (EICEs)^13^, while interactions between IRF8 and AP-1 drive transcriptional activity from AP-1-IRF combinatorial elements (AICEs)^14^. IRF2 is hypothesized to be a transcriptional repressor which acts to inhibit IRF1 and IRF9^15,16^, though IRF2 has also been shown to act as a transcriptional activator for select proteins^17,18^. IRF2 has been identified to bind a number of proteins, including PU.1^19^, IRF1^15^, and IRF8^20^, though the physiological significance of these interactions is not currently understood.

Loss of IRF2 within mice has pleiotropic effects. IRF2 is an inhibitor of the IFNAR signaling pathway, and *Irf2*^*-/-*^ mice develop an IFNAR-dependent CD8 T cell mediated inflammatory skin disorder^16^. *Irf2*^*-/-*^ mice also develop IFNAR-dependent progressive anemia characterized by a loss of late-stage erythroblasts within the bone marrow^21^. IRF2 also functions as a tumor suppressor, and loss of IRF2 activity has been found in a number of human primary tumors^22^. *Irf2*^*-/-*^ mice experience a significant loss of splenic DC2s, as measured by expression of the DC2 markers CD4 and CD11b^1,2^. However, Flt3L culture of *IRF2*^*-/-*^ bone marrow fails to generate either DC1s or DC2s-a surprising result given that DC1 development was fully intact in the spleens of *IRF2*^*-/-*^ mice^1,2^. *Irf2*^*-/-*^ mice additionally experience a loss of mature CD11b^+^ NK cells and an expansion of basophils in the spleen, liver, and blood^23,24^. Loss of CD11b^+^ splenic DC2s was dependent on IFNAR, and *Irf2*^*-/-*^ *Ifnar*^*-/-*^ mice exhibited normal numbers of splenic DC2s^1,2^. However, neither loss of NK cells nor expansion of basophils is dependent on expression of IFNAR, suggesting involvement of IRF2 in multiple developmental pathways^1^.

*Irf2*^*-/-*^ mice also develop acute pancreatitis on exposure to polyI:C, possibly mediated by a failure of IRF2 to suppress IRF5- and IRF7 mediated *trypsinogen5* expression^25^.

The role of IRF2 within DC2 development remains uncertain. The original finding that *Irf2*^*-/-*^ mice lacked a significant majority of their splenic CD4^+^ CD11b^+^ DC2s has not been followed up on. While the authors performed a bone marrow chimera to determine that the requirement for IRF2 in DC2 development was hematopoietic-intrinsic, their assay was unable to determine whether the requirement for IRF2 was cell-intrinsic^1^. Here, we show that IRF2 is not required in a cell-intrinsic manner for cDC2 development. We additionally demonstrate that loss of IRF2 or addition of IFNα is detrimental to cDC development in Flt3L-treated bone marrow cultures, and that simultaneous loss of IRF2 and IFNAR restores cDC development from Flt3L-treated bone marrow. We propose that loss of cDC2s in *Irf2*^*-/-*^ mice is caused by an exogenous effect of IFNα on cDC2 development.

## Materials and Methods

### Mice

WT C57BL6/J and *Irf2*^−*/*–^ (B6.129S2-Irf2^*tm1Mak*^/J) mice were obtained from the Jackson Laboratory. B6.SJL (B6.SJL-*Ptprc*^*a*^ *Pepc*^*b*^ /BoyJ) mice were obtained from Charles River. All mice were maintained on the C57BL/6 background in the Washington University in St. Louis School of Medicine specific pathogen-free animal facility following institutional guidelines with protocols approved by Animal Studies Committee at Washington University in St. Louis. Experiments were performed with mice between 6 and 10 weeks of age unless otherwise noted.

### Dendritic cell preparation

Splenic DCs were harvested and prepared as described previously^26^. Briefly, spleens and were minced and digested in 5 mL of Iscove’s modified Dulbecco’s media (IMDM) + 10% fetal calf serum (FCS) (cIMDM) with 250 μg/mL collagenase B (Roche) and 30 U/mL DNaseI (Sigma-Aldrich) for 45 min at 37ºC with stirring. After digestion was complete, single cell suspensions from all organs were passed through 70-μm strainers and red blood cells were lysed with ammonium chloride-potassium bicarbonate (ACK) lysis buffer. Cells were subsequently counted with a Vi-CELL analyzer (Beckman Coulter), and 3-5×10^6^ cells were used per antibody staining reaction.

### Flow cytometry

Cells were kept at 4ºC while being stained in PBS supplemented with 0.5% BSA and 2mM EDTA in the presence of antibody blocking CD16/32 (clone 2.4G2; BD 553142). All antibodies were used at a 1:200 dilution vol/vol (v/v), unless otherwise indicated. The following antibodies were used: V500–anti-MHC-II (clone M5/114.15.2), Alexa Fluor 700 or peridinin chlorophyll protein (PerCP)-eFlour 710–anti-CD11b (clone M1/70), Brilliant Violet 510 or PE/Dazzle 594– anti-CD45R (clone RA3-6B2), allophycocyanin(APC)–anti-CD317 (clone eBio927, 1:100 v/v), PE-Cy7–anti-CD24 (clone M1/69), PerCP–eFluor 710–anti-CD172a (clone P84), v450–anti-CD8α (clone 53-6.7), Alexa Flour 700 or APC/Cy7–anti-F4/80 (clone BM8, 1:100 v/v), Brilliant Violet 605 or PE–anti-CD45.2 (clone 104), Brilliant Violent 650–anti-CD45.1 (clone A20), and APC–eFluor 780–anti-CD11c (clone N418). Cells were analyzed on a FACSAria Fusion flow cytometer (BD), or an Aurora spectral flow cytometer (Cytek), and data were analyzed with FlowJo v10 software (TreeStar).

### Bone marrow chimeras

CD45.1^+^ mice (B6.SJL-*Ptprc*^*a*^ *Pepc*^*b*^ /BoyJ) were lethally irradiated with a dose of 1050 rads. 24 hours later mice were injected i.v. with either CD45.2^+^ wild-type BM, CD45.2^+^ *Irf2*^−*/*–^ bone marrow, or a 90:10 mixture of CD45.2^+^ *Irf2*^−*/*–^ and CD45.1^+^ wild-type bone marrow. Mice were analyzed 20 weeks later.

### Cell culture

Bone marrow was isolated from mouse femurs, tibias, and hips. Bones were crushed in phosphate-buffered saline supplemented with 5% FCS and passed through 70-μm strainers to create single-cell suspensions. Red blood cells were lysed using ACK lysis buffer. Cells were then counted with a Vi-CELL analyzer (Beckman Coulter) and plated at a density of 8×10^6^ cells per well in 6-well plates. In indicated experiments, interferon α4 (PBL Assay Science) was added to cell cultures. Cells were cultured in cIMDM supplemented with Flt3L-conditioned media for 7 day at 37C prior to analysis.

### Statistics

All statistical analyses were performed using GraphPad Prism software version 8. Unless otherwise noted, Mann-Whitney test was used to determine significant differences between samples. Values are presented as the average ± SEM. P≤0.05 was considered statistically significant.

## Results

### IRF2 is required in a cell-extrinsic manner for cDC2 development

To verify that cDC2s were dependent of IRF2 for their development, we obtained *Irf2*^*-/-*^ mice and analyzed their spleens. We observed a loss of cDC2s in the spleens of *Irf2*^*-/-*^ mice relative to wild-type mice, confirming previous observations (Fig. 1A, 2A). We next set to test whether IRF2 was required for the development of the ESAM^+^ subset of cDC2s, similar to Notch2 and RelB^5,8^. We observed no difference in ESAM expression within remaining cDC2s within *Irf2*^*-/-*^ mice, confirming that IRF2 is not required for the development of Notch2-dependent cDC2s (Fig. 1B).

**Figure 1:**
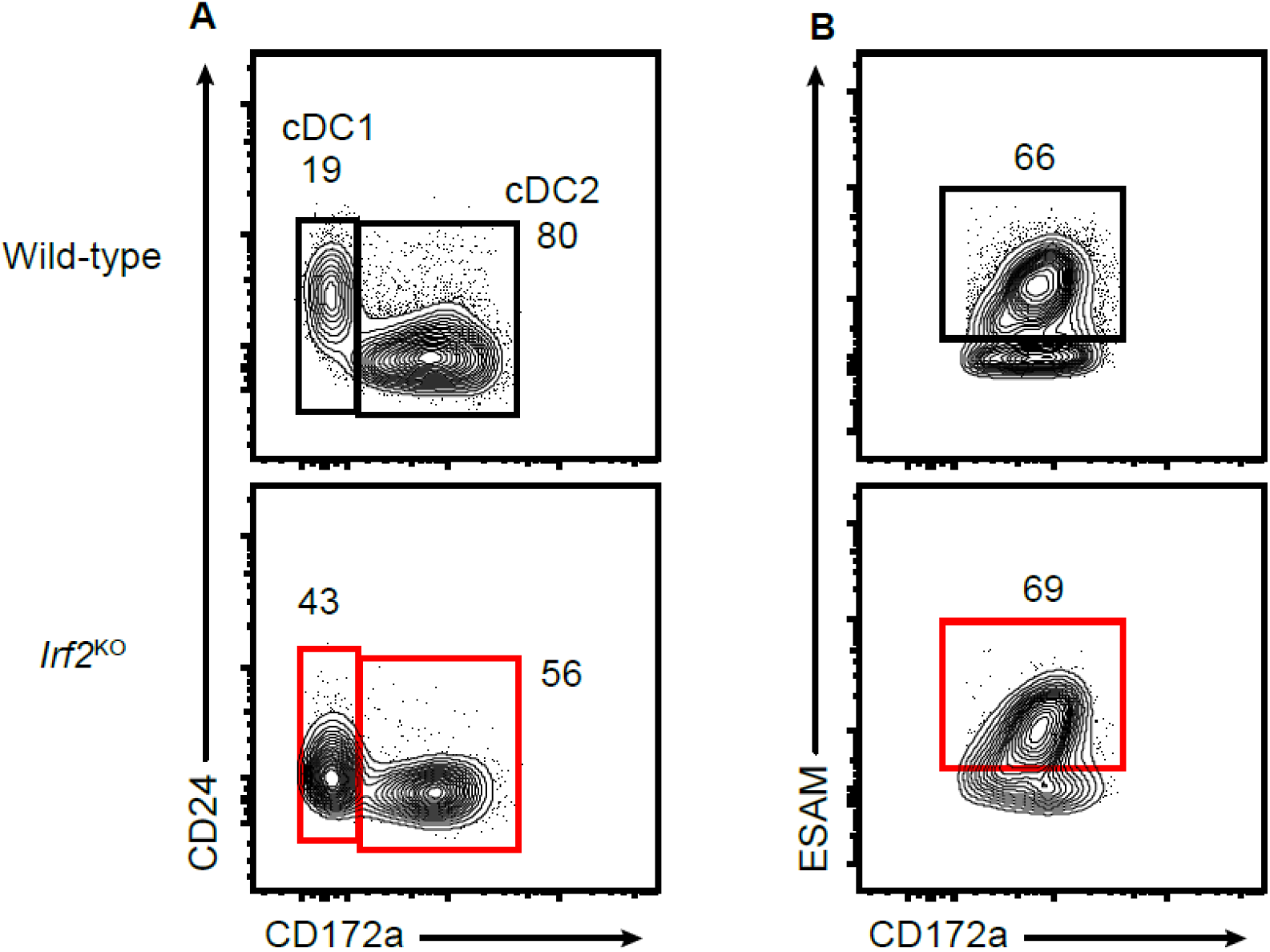
Reduced cDC2 in *Irf2*^*-/-*^ mice. **(A)** Spleens were analyzed from wild-type or *Irf2*^−*/*–^ (*Irf2*^KO^) mice for cDC1 and cDC2. Shown analysis is pregated on CD317^-^ CD45R^-^ MHCII^+^ CD11c^+^ single cells. **(B)** cDC2 from wild-type or *Irf2*^*KO*^ spleens were analyzed for expression of the surface marker ESAM. Shown analysis is pregated on CD317^-^ CD45R^-^ MHCII^+^ CD11c^+^ CD24^-^ CD172α^+^ single cells.

*Irf2*^*-/-*^ mice acquire progressive dermatitis as they age^16^. To determine whether there was a relationship between acquisition of dermatitis and loss of cDC2s, we evaluated whether cDC2s were progressively lost in aging mice. We found that *Irf2*^*-/-*^ mice had fewer splenic cDC2s as a fraction of total cDCs at 15-20 weeks of age than they did at 6-10 weeks of age (Fig 2B), suggesting that loss of cDC2s in *Irf2*^*-/-*^ mice might be cell-extrinsic and related to inflammation within these mice. To test whether IRF2 was required in a cell-intrinsic manner in cDC2s, we set up mixed bone marrow chimeras, transplanting a 50:50 mixture of either CD45.1^+^ wild-type and CD45.2^+^ wild-type bone marrow or CD45.1^+^ wild-type and CD45.2^+^ *Irf2*^*-/-*^ bone marrow into lethally-irradiated wild-type mice. We found that *Irf2*^*-/-*^ bone marrow competed poorly with wild-type bone marrow for engraftment, as previously reported^27^. However, we observed that *Irf2*^*-/-*^ bone marrow was able to generate pDCs, cDC1s, and cDC2s at similar rates, indicating no specific defect in formation of cDC2s from *Irf2*^*-/-*^ bone marrow (Fig. 3A, B). We therefore conclude that IRF2 is required for cDC2 development in a cell-extrinsic manner.

**Figure 2:**
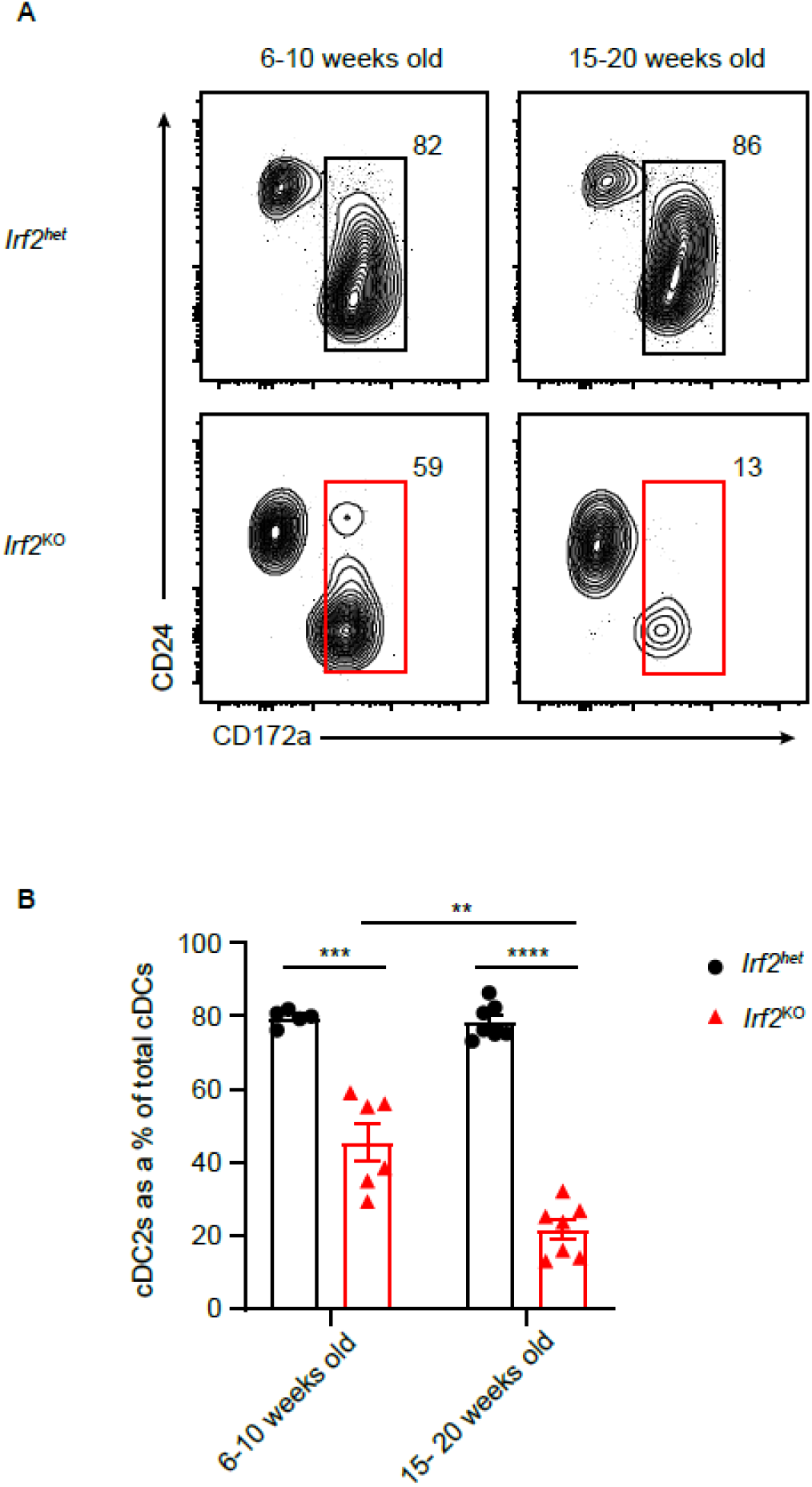
Magnitude of cDC2 loss in *Irf2*^*-/-*^ mice increases with age. **(A)** Spleens were analyzed from wild-type or *Irf2*^−*/*–^ (*Irf2*^KO^) mice that ranged from 6 to 10 weeks old or 15 to 20 weeks old for cDC2. Shown analysis is pregated on CD317^-^ CD45R^-^ MHCII^+^ CD11c^+^ single cells. **(B)** Quantification of cDC2 (gated as CD11c^+^ MHCII^+^ CD24^-^ CD172α^+^ single cells) as a function of total cDCs within the spleen. ** *p* < 0.01, *** *p* < 0.001, **** *p* < 0.0001 (Student’s *t*-test).

**Figure 3:**
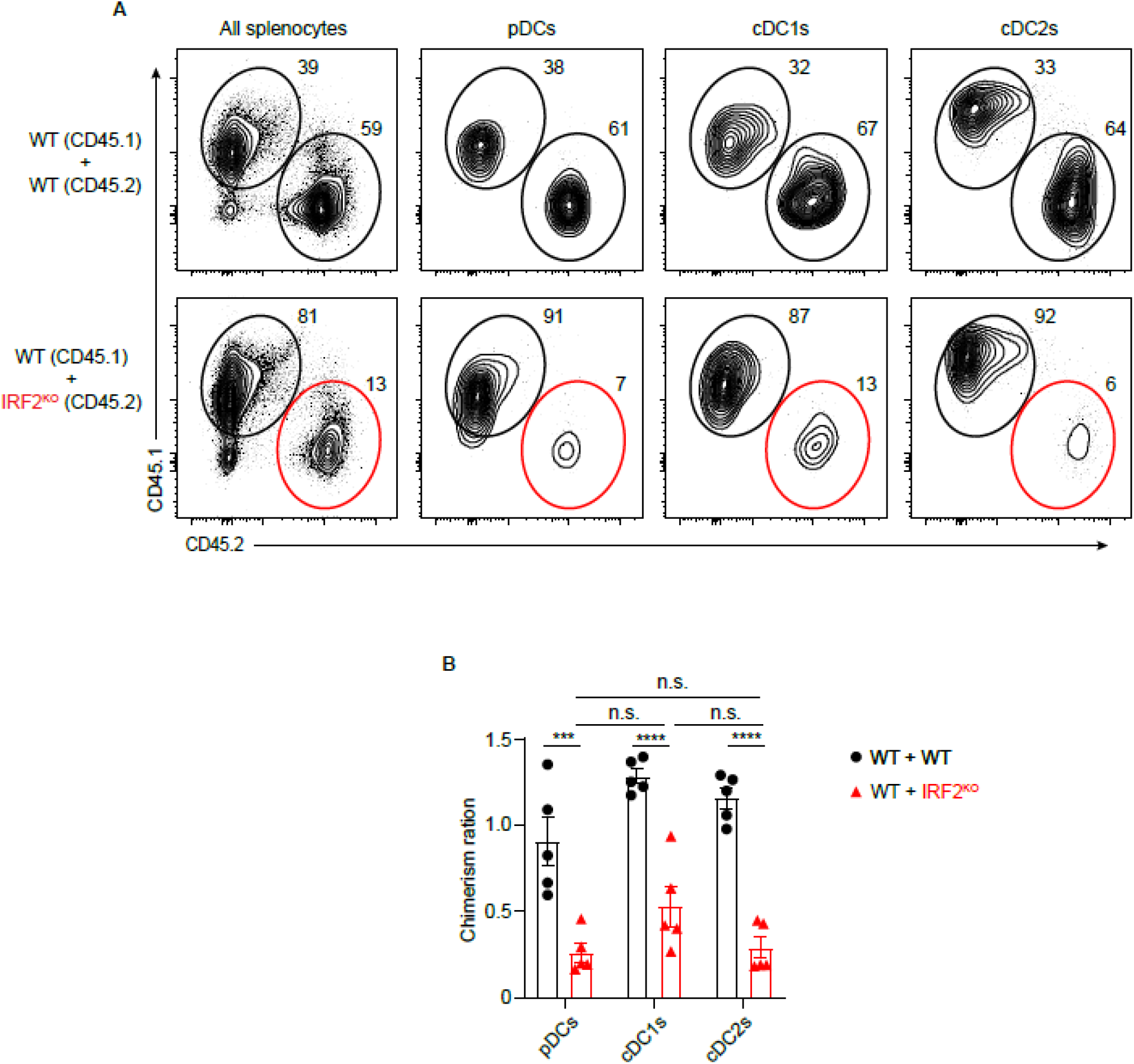
Effects of *Irf2* on cDC2 formation are not cell-intrinsic. **(A)** FACS analysis of spleens from chimeras generated with equal mixes of CD45.1^+^ B6.SJL and CD45.2^+^ wild-type (WT) or CD45.2^+^ *Irf2*^-/-^ (IRF2^KO^) bone marrow analyzed 20 wk after lethal radiation and transplant. **(B)** Contributions of CD45.2^+^ bone marrow to the indicated lineages. Chimerism ratio calculated as the proportion of the indicated lineage that is CD45.2^+^ divided by the proportion of all lymphocytes that are CD45.2^+^. *** *p* < 0.001, **** *p* < 0.0001, n.s. not significant (Student’s *t*-test).

### Loss of IRF2 or addition of IFNα inhibits cDC cultures in an IFNAR-dependent manner

To evaluate the effects of loss of IRF2 on cDC development, we turned to in vitro culture systems. We cultured wild-type or *Irf2*^*-/-*^ with Flt3L for 7 days. We observed a modest reduction in CD45R^+^ CD317^+^ pDCs, though it was uncertain whether this was due to a genuine loss of pDCs or merely loss of the CD45R surface marker on pDCs (Fig. 4A). We also observed an increase in expression of the interferon-stimulated gene CD317 (Fig. 4A). Notably, we also found that while wild-type bone marrow was able to give rise to a robust population of MHCII^+^ CD11c^+^ cDCs, *Irf2*^*-/-*^ bone marrow was incapable of giving rise to any such population (Fig. 4B). Zbtb46 is a transcription factor which marks conventional dendritic cells^28^. As IRF2 is an antagonist of type 1 IFN signaling, we treated *Zbtb46*^*gfp/+*^ bone marrow with Flt3L and varying doses of IFNα. While pDC numbers were not substantially changed (Fig. 5A), we noticed a striking decrease in Zbtb46-GFP^+^ cDCs in IFNα-treated cultures (Fig. 5B, C). Higher doses of IFNα resulted in an almost complete loss of cDC development, reminiscent of cultured *Irf2*^*-/-*^ bone marrow (Fig. 4B, 5B).

**Figure 4:**
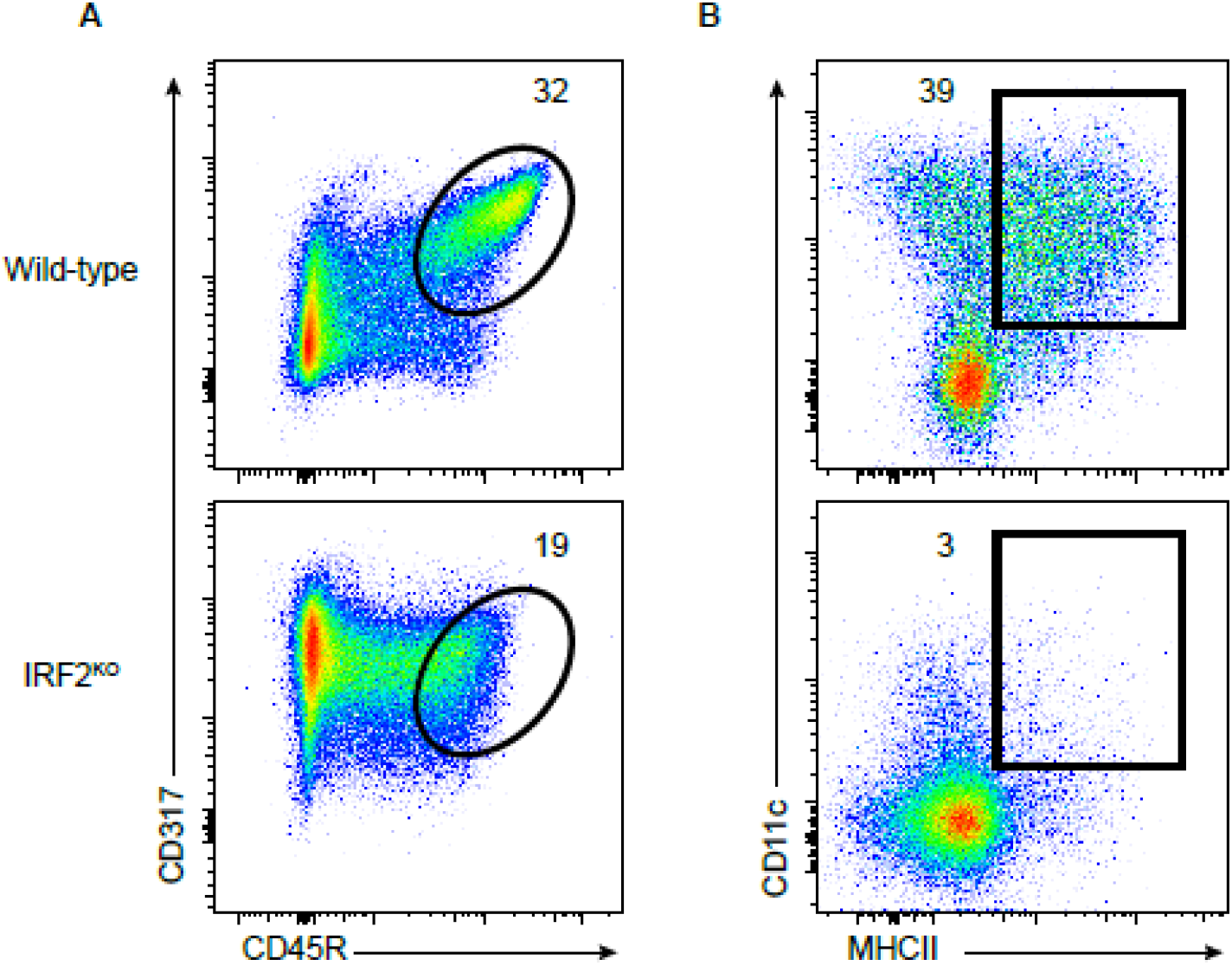
Defective formation of cDCs in cultured *Irf2*^*-/-*^ bone marrow. **(A)** FACS analysis of wild-type or IRF2^KO^ bone marrow cultures treated with Flt3L for 7 days for development of CD317^+^ CD45R^+^ pDCs. Cells pregated on single cells. **(B)** FACS analysis of wild-type or IRF2^KO^ bone marrow cultures treated with Flt3L for development of CD11c^+^ MHCII^+^ cDCs. Cells pregated on CD45R^-^ single cells.

**Figure 5:**
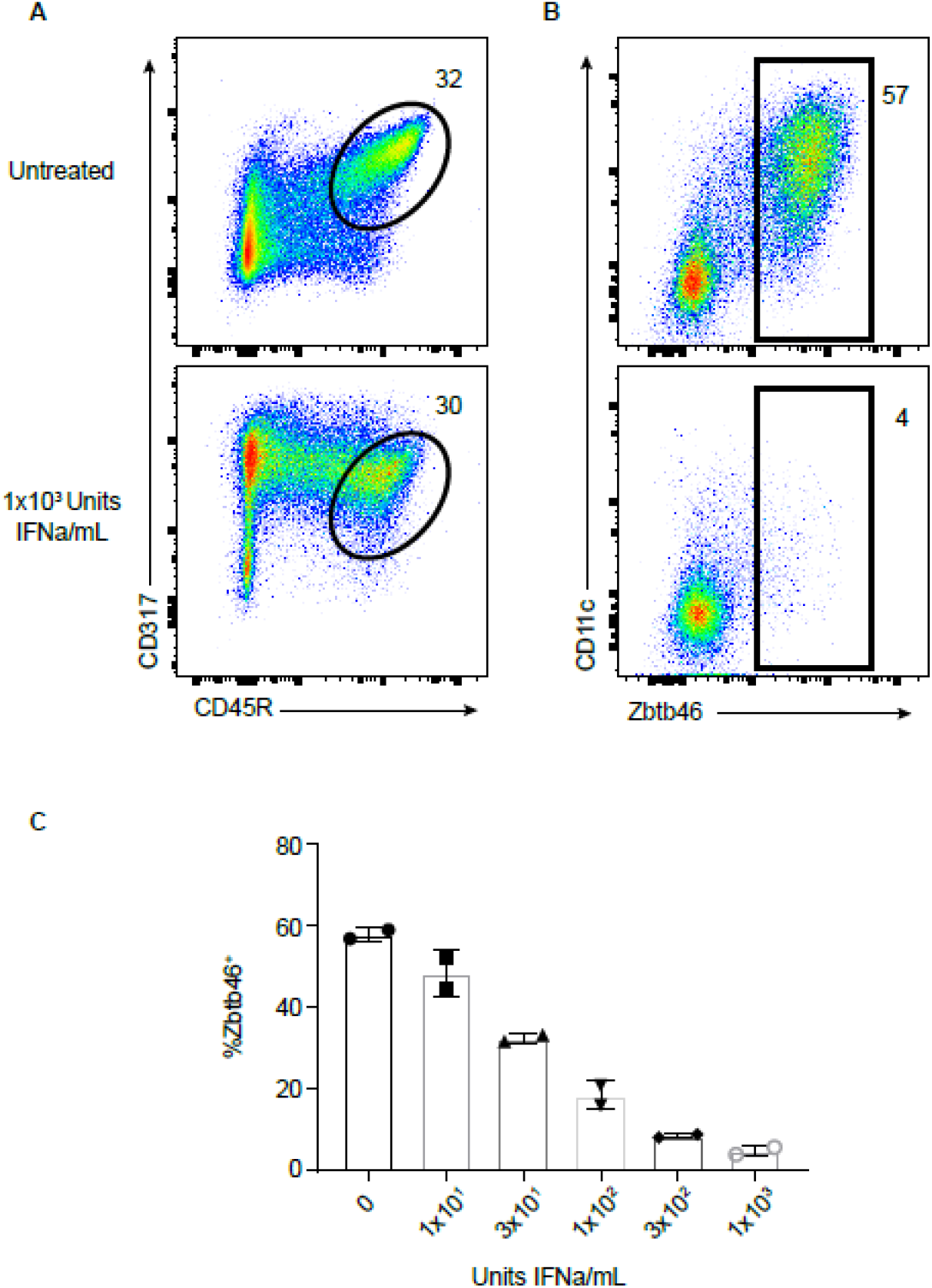
IFN-α treatment of Flt3L cultures abrogates cDC development. **(A)** FACS analysis of Zbtb46^GFP/+^ bone marrow cultures treated with Flt3L with or without 10^3^ Units/mL IFNα for 7 days for development of CD317^+^ CD45R^+^ pDCs. Cells pregated on single cells. **(B)** FACS analysis of Zbtb46^GFP/+^ bone marrow cultures treated with Flt3L with or without 10^3^ Units/mL IFNα for 7 days for development of Zbtb46^+^ cDCs. Cells pregated on CD45R^-^ single cells. **(C)** Analysis of Zbtb46^GFP/+^ bone marrow cultured for 7 days with Flt3L and varying quantities of IFNα as indicated. Presented are Zbtb46^+^ cDCs as a proportion of CD45R^-^ single cells.

To test specifically whether IFNα was responsible for phenotypes induced by loss of IRF2, we crossed *Ifnar1*^*-/-*^ mice lacking the type 1 IFN receptor to *Irf2*^*-/-*^ mice to generate *Irf2*^*-/-*^*Ifnar1*^*-/-*^ mice. These mice, though lacking IRF2, are insensitive to the effects of IFNα signaling, and do not develop the dermatitis that characterizes *Irf2*^*-/-*^ mice^16^. We found that *Irf2*^*-/-*^ *Ifnar1*^*-/-*^ mice generated splenic cDC2 numbers which did not significantly differ from normal (Fig. 6A, B), confirming previous reports^1^. Additionally, we found *Irf2*^*-/-*^ *Ifnar1*^*-/-*^ mice developed cDCs as normal (Fig. 6C). This establishes that loss of cDCs in Flt3L-cultured *Irf2*^*-/-*^ bone marrow is due to effects of type 1 interferons.

**Figure 6:**
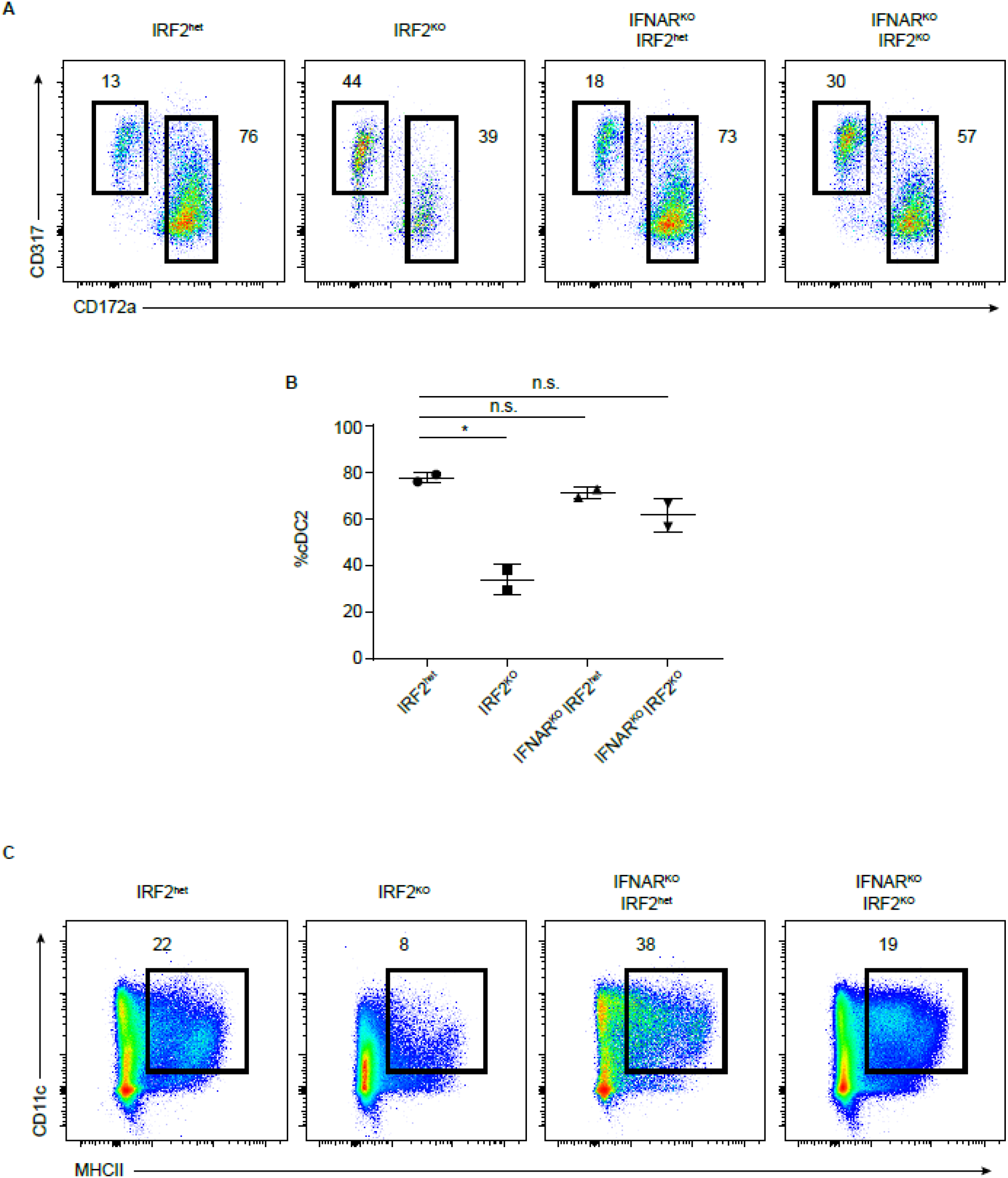
*Ifnar* signaling mediates defective DC development in *Irf2*^*-/-*^ mice. **(A)** Spleens from mice of the indicated genotypes were analyzed for cDC1 and cDC2. Shown analysis is pregated on CD317^-^ CD45R^-^ MHCII^+^ CD11c^+^ single cells. **(B)** Quantification of cDC2 (gated as CD11c^+^ MHCII^+^ CD24^-^ CD172α^+^ single cells) as a function of total cDCs within the spleen. **(C)** FACS analysis of bone marrow cultures of the indicated genotypes treated with Flt3L for 7 days for development of CD11c^+^ MHCII^+^ cDCs. Cells pregated on CD45R^-^ single cells. * *p* < 0.05, n.s. not significant (Student’s *t*-test).

## Discussion

Here, we establish that IRF2 is required for normal development of cDC2s, but that this requirement is cell-extrinsic. IRF2 was first identified in 1989 as a gene which bound similar genomic sequences to IRF1, and which was able to inhibit IRF1 activity by outcompeting IRF1 in binding to these sequences^15^. Specific functions and mechanisms of IRF2 activity are still unclear. IRF2 has been reported both as a transcriptional activator and as a transcriptional repressor^15,17,29^, and its mechanism of action has not been fully established. IRF2 is hypothesized to inhibit signaling through the IFNα receptor by blocking the ability of the downstream ISGF3 complex composed of IRF9, STAT1, and STAT2 to associate with interferon-stimulated response element sequences termed ISREs^16^. However, IRF2 is also known to have effects outside of the type 1 interferon pathway. *Irf2*^*-/-*^ mice show increased numbers of basophils which are not restored to normal numbers by loss of IFNAR1^24^, in contrast to our findings in DCs (Fig. 6). We and others observe dysregulated IFNα activity in *Irf2*^*-/-*^ mice, but it remains unclear in which cell types IRF2 acts to control IFNα activity. IRF2 has recently been proposed regulate human keratinocyte development and murine intestinal stem cell development, raising the possibility that IRF2 acts in non-immune lineages to exert control over the immune system^30,31^. Recent development of a conditionally-deleted IRF2 allele should greatly facilitate efforts to determine specific cell types that IRF2 acts in which might influence cDC2 development^31^.

Strikingly, development of cDC2s, but not cDC1s, is reduced in *Irf2*^*-/-*^ mice. While recent work from our lab has thoroughly characterized steps and factors involved in cDC1 development^26,32^, cDC2 development remains opaque. The mechanism by which cDC2s are lost in *Irf2*^*-/-*^ mice is not immediately obvious. Developing cDC2s could be diverted into pDCs or cDC1s by signaling through IFNAR1. Another hypothesis is that IFNAR1 signaling is relatively toxic to cDC2s than cDC1s, resulting in the preferential loss of cDC2s in vivo. From a population perspective, it may be evolutionarily beneficial to reduce cDC2 development and favor cDC1 development in response to IFNα signaling. cDC1s are capable of cross-presenting antigen to activate cytotoxic CD8^+^ T cells^33^. Induction of type 1 IFN responses to viruses may support cDC1 development by blocking cDC2 development, in turn increasing the number of cDC1s able to support CD8^+^ T cell activation.

We found that IFNα treatment of Flt3L bone marrow cultures induced loss of both cDC1s and cDC2s, but not pDCs (Fig. 5). Similarly, Flt3L bone marrow cultures created using *Irf2*^*-/-*^ bone marrow notably lack cDC1s and cDC2s (Fig. 6). *Irf2*^*-/-*^ cDC1s are therefore able to develop in vivo but not in vitro, reminiscent of our observations of *Bcl6*^*cKO*^ cDC1s (Chapter 1)^34^.

Conversely, NFIL3 is known to be required for cDC1 development in vivo^35^ but not in vitro^32^. Future work will need to investigate transcriptional pathways governing cDC1 differentiation in vitro to determine how these pathways differ from in vivo developmental pathways.

Further investigation into cDC2s will be necessary to characterize the transcriptional machinery governing these cells’ development. It has been demonstrated that a monocyte-dendritic cell precursor (termed the MDP) gives rise to a common dendritic precursor (termed the CDP), which in turn gives rise to a pre-cDC2 that represent the immediate progenitor population^36-40^. Our lab has recently demonstrated that commitment to cDC1 development initiates as early as the CDP with the activation of NFIL3 expression in a population of CDPs^32^. It is currently uncertain at which stage progenitors commit to cDC2 development. cDC1 progenitors which are unable to autoactivate IRF8 expression fail to develop into mature cDC1s and instead will develop into cDC2s, raising the possibility that cDC2s in vivo are “developmentally failed” cDC1s^39^. However, we have recently found that mutation of *Zeb2* enhancer elements selectively induced loss of cDC2 development^4^. Further, simultaneous mutation of *Zeb2* and *Irf8* enhancer elements restores mature cDC2s, but not pre-cDC2s, using a “developmentally failed” cDC1 route^4^. This demonstrates that “developmentally failed” cDC1s are not a major source of cDC2s in wild-type mice. We therefore hypothesize that cDC2 development uses distinct transcriptional machinery from cDC1 development. Work in this thesis demonstrates that IRF2 is not a cell-intrinsic part of this machinery, but that type 1 interferon signaling is able to affect cDC2 development through the IFNα receptor. Future work in this field will be needed to elucidate exact mechanisms of cDC2 development, as well as to determine how these mechanisms are perturbed in *Irf2*^*-/-*^ mice.

